# Investigating the genetic architecture of biotic stress response in stone fruit tree orchards under natural infections with a multi-environment GWAS approach

**DOI:** 10.1101/2024.10.15.618428

**Authors:** Marie Serrie, Vincent Segura, Alain Blanc, Laurent Brun, Naïma Dlalah, Frédéric Gilles, Laure Heurtevin, Mathilde Le-Pans, Véronique Signoret, Sabrina Viret, Jean-Marc Audergon, Bénédicte Quilot, Morgane Roth

**Affiliations:** INRAE, UR GAFL, Avignon, France; AGAP Institut, Université Montpellier, CIRAD, INRAE, Institut Agro, Montpellier, France; Geno-Vigne®, IFV, INRAE, Institut Agro, Montpellier, France; INRAE, UE A2M, Avignon, France; INRAE, UERI Gotheron, Saint-Marcel-Lès-Valence, France

**Keywords:** Biotic stress, genome-wide association studies (GWAS), multi-environment trials (METs), Genotype × Environment interactions, resistance, tolerance, stone fruit tree, low phytosanitary inputs, core collections, *Prunus*

## Abstract

The mapping and introduction of sustainable plant immunity to pests and diseases in fruit tree is still a major challenge in modern breeding. This study aims at deciphering the genetic architecture underlying resistance or tolerance across environments for major pests and diseases in peach (*P. persica*) and apricot (*P. armeniaca*). We set up a multi-environment trial (MET) approach by studying two core collections of 206 peach and 150 apricot accessions deployed under low phytosanitary conditions in respectively three and two environmentally contrasted locations in South-East of France. To capture the complex dynamics of pest and disease spread in naturally infected orchards, visual scoring of symptoms was repeated within and between 3 years, for five and two pests and diseases respectively for peach and apricot, resulting in the maximum of damage score and the AUDPC. These traits were used as phenotypic inputs in our genome-wide association studies (GWAS) strategy, and leading to the identification of: i) non-additive genotype–phenotype associations, ii) environment-shared QTLs iii) environment-specific QTLs, and iv) interactive QTLs which changes in direction (‘antagonist’) or intensity (‘differential’) according to the environment. By conducting GWAS with multiple methods, we successfully identified a total of 60 high confidence QTLs, leading to the identification of 87 candidate genes, the majority belonging to the Leucine-rich repeat containing receptors (LRR-CRs) family gene. Finally, we provided a comparative analysis of our results on peach and apricot, two closely related species. The present results contribute to the development of genomics-assisted breeding to improve biotic resilience in *Prunus* varieties.

## Introduction

Biotic stresses represent a major threat to agricultural production with a massive diversity of organisms being able to affect plants, including a large range of viruses, bacteria, fungi, phytoplasma, nematodes and pests [1,2]. Reducing the impact of these biotic stresses in a sustainable and environment-friendly way is urgent and, as such, reducing plant susceptibility through genetic progress appears as an effective and promising strategy [3]. It has been long recognized that host defence against parasites and pathogens can be divided into two conceptually different mechanisms: the ability to reduce and control pathogen (resistance) and the ability to limit the damage caused by a given within-host pathogen load without necessarily reducing this pathogen load (tolerance) [4,5]. Developing cultivars that present durable low susceptibility to biotic stresses is therefore an important breeding goal, in particular for perennial woody crops, given their long generation times and lifespans in orchards [6]. After multiple episodes of resistance breakdown due to the introgression into crop cultivars of monogenic resistances [7,8], research is now focusing on polygenic defence traits which are likely more durable but also more complex to understand and to introduce in elite lines [9]. The combination of major resistance genes with quantitative resistance/tolerance factors have already shown promising results as an alternative to ensure greater durability of crop protection [10,11]. In addition to specific host resistances genes, broad-spectrum defence factors against multiple diseases represent another promising solution for sustainable plant defence [12]. Some degree of tolerance has already been observed in different landraces cultivar and wild relatives, as *P*. *davidiana*, a relative of peach (*P. persica*) carrying resistance genes against green peach aphids, powdery mildew and *Plum pox virus* (PPV) [13–15]. In this context, the natural genetic diversity of crops is a valuable source of untapped favourable allelic combinations which deserves to be explored [16].

Genome Wide Association Studies (GWAS) have been extensively used to understand the genetic basis of complex traits such as quantitative disease resistance or tolerance [17]. This whole-genome approach exploits ancestral linkage disequilibrium among diverse germplasm to detect associations between phenotypic traits and polymorphisms, also called quantitative trait loci (QTL) [18,19]. QTLs identified by GWAS can be used in breeding programs through the development of marker-assisted selection (MAS) in relevant germplasm [20,21]. Yet, detecting QTLs related to disease resistance or tolerance can be challenging since phenotypic variation is often modulated by environmental effects and interactions between genotype and environment (G×E) [22–24]. The consequences of G×E can be illustrated by a change in the phenotypic ranking of individuals across environments which could be explained by a modulation of the size and/or sign of the QTL effect by the environment. In this context, Multi-Environment Trials (MET), consisting in phenotyping the same panel of genotypes in different locations and/or over different years, have been widely adopted to describe QTL responses to different environmental factors [25,26]. To detect QTLs of interest from METs, a first approach is to perform single-environment GWAS to identify and further compare QTLs between specific environments [27,28]. A complementary approach is based on multi-environment GWAS in order to detect QTLs with both main effects and QTL × Environments effects (*i.e.* QTLs with differential effects depending on environmental conditions) [27]. Models have been developed to handle QTL × Environments interactions, such as the multi-trait mixed model MTMM [29] which has been successfully used, firstly in *Arabidopsis thaliana* for the study of flowering time QTLs under different temperatures, and later for plant architecture in barley and in production traits in common bean under contrasted abiotic stress conditions [30–32]. In addition, single-environment GWAS can be exploited via meta-analysis GWAS to identify QTLs with minor effects which might go undetected in separate analyses, as well as to harness markers with stable effects across environments [33,34]. Meta-GWAS are now emerging in plant studies as a practical alternative for combining multiple independent studies to identify additional marker-trait associations and to verify the association previously identified [33,35,36]. A useful aspect of Meta-GWAS for plant is that it can be applied on unbalanced phenotypic datasets (*i.e.* asymmetrical proportions between class of observation) and it does not require replication of the same individuals across all environments [37].

Taking advantage of the recent publication of reference genomes [38–41] and of the availability of molecular technology such as SNP array [42], GWAS has gradually become more widespread in *Prunus* species for determining the genetic basis of variation in complex agronomic or fruit quality traits [43–47]. Despite these methodological advances, to date G×E interaction effects and the genetic determinants of partial resistance to different biotic stresses from the wide diversity in *Prunus* fruit trees has been little investigated [48–53]. One GWAS experiment has explored non-additive effect in almond and allow the identification of 13 QTLs for several agronomical traits, with only one having additive effects [54]. These results underline that in heterozygous tree crops a significant share of non-additive variance can be exploited in GWAS. Statistical estimates of non-additive effects can therefore allow for a more accurate prediction of total genetic variance which results in a more complete characterization of the genetic architecture of disease resistance traits and help to find complementary QTLs of interest [55,56].

In this context, our objective is to dissect the genetic architecture of pests and diseases resistance or tolerance in peach (*Prunus persica* L.) and apricot (*Prunus armeniaca* L.) in multiple environments by using GWAS. GWAS analysis are particularly well suited for studying peach and apricot genomes by i) being diploid with small size genomes (with respectively 230 and 295 Mbp) [57], ii) harbouring a large untapped phenotypic diversity available in germplasm collection [58,59], and iii) being highly syntenic, thus allowing comparison between them [60]. We built a comprehensive database gathering the phenotypic response to biotic stresses of peach and apricot using core collections of respectively 206 and 150 accessions monitored under low phytosanitary protection in three and two different orchards in the South-East of France. The pests and diseases observed included leaf curl, leafhopper, powdery mildew, rust and shot hole for peach as well as blossom blight and rust for apricot. We conducted a series of complementary GWAS to i) firstly investigate the main additive and non-additive genetic effects across environments, secondly ii) compare environment specific effects with single-environment GWAS, and finally iii) harness G×E interactions effects using multi-environment GWAS and meta-analysis GWAS.

## Results

### Phenotypic response of peach and apricot accessions to natural infection in multiple orchards

The first step of our analyses aimed at evaluating the biotic performance of the two core collections and to assess the effects of environment, genotype and genotype by environment interactions on biotic stress responses.

### A large variability in the phenotypic response to biotic stress in the two core collections

We scored trees at different time points for each biotic stress throughout the season to fully capture the dynamic expression of the phenotypic diversity under low phytosanitary protection. This allowed us to produce two types of variables for each tree and each biotic stress, namely the i) maximum of susceptibility (Max) and ii) the Area under the Disease Progress Curve (AUDPC).

After spatial correction, we cleaned our dataset in order to retain only environments with sufficient expression of phenotypic variability for each trait, leading to the removal 19.6% of the data (Supplementary Method 2 and Supplementary Fig 1). In the remaining dataset the average standard deviation was of 31.1 and 295.4 respectively for Max and AUDPC (Figure 1 and Supplementary Fig 2). We found the highest diversity of response for apricot rust leaf fall for both Max and AUDPC (with standard deviation of 34.1 and 321.9 respectively). For an equivalent value of Max, trees presented markedly different temporal trajectories for this disease, as illustrated by diverging onset dates and spread of symptoms, with marked environment-specific effects (Figure 1 and Supplementary Fig 3). While correlations between Max and AUDPC remained high for all biotic stresses (from 0.676 to 0.997) (Figure 1 and Supplementary Fig 2), AUDPC thus provided complementary information on the response of accessions to biotic stresses.

**Figure 1.**
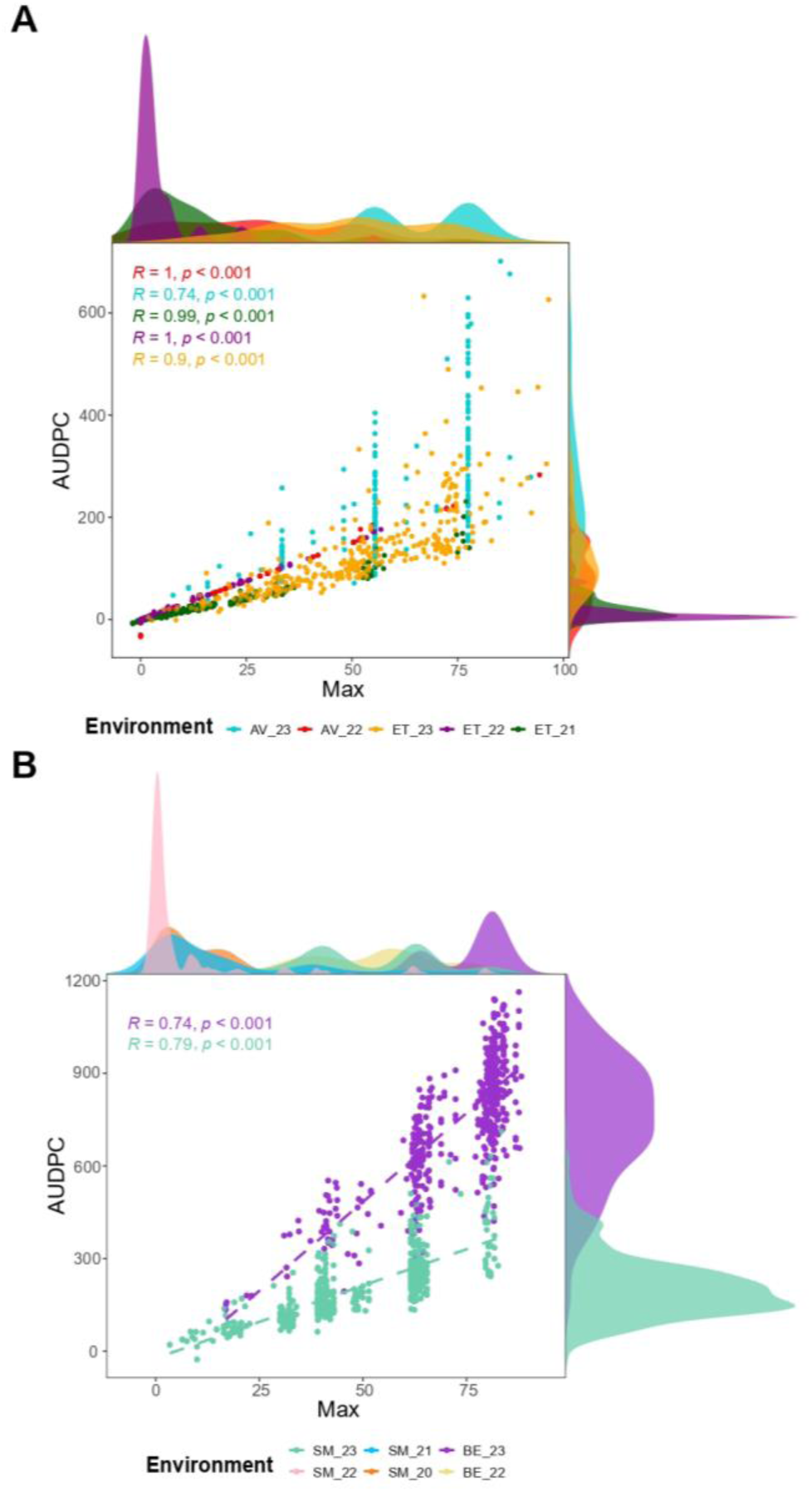
Phenotypic distribution and correlation between Max and AUDPC for A) peach rust and B) apricot rust. Distributions of Max are shown at the top of each graph, while the distributions of AUDPC are shown on the right. *AV: Avignon, ET: Étoile-sur-Rhône, TO: Torreilles, SM: Saint-Marcel-lès-Valence and BE: Bellegarde*.

To characterize the stability of symptom expression across environment, we calculated the correlations between environment-specific BLUPs for each pest or disease. The genetic correlations between environments ranged from moderate to strong, with average Pearson’s correlation coefficients of 0.68 and 0.73 for Max and AUDPC respectively (Supplementary Table 1 and Supplementary Fig 4). We found the lowest genetic correlation between environments for blossom blight (*r* = 0.41 in average for Max) and the highest correlations for leaf curl (*r* = 0.87 and *r*= 0.91 in average for Max and AUDPC respectively) (Supplementary Table 1). Overall, symptoms were more correlated within a given location than within a given year for both Max and AUDPC (Supplementary Table 1).

### Biotic stress responses are modulated by G and G×E effects

We computed a linear mixed model for each pest and disease to quantify the variance associated to genotype, genotype by location, genotype by year and genotype by year by location interactions (Figure 2.A and Supplementary Method 3). The variance attributed to the genotype amounted between 5.1 and 66.7 % of the total variance. For powdery mildew and shot hole, the genotypic variance was lower than the variance attributed to the G×L interactions (for both Max and AUDPC variables), and for blossom blight lower than the variance attributed to the G×Y interactions (39.7% of total variance). The contribution on the G×L effect on the trait variance was moderate to high (6.3– 26.5%) and was higher than the variance explained by G×Y for all biotic stresses, as already expected from previous correlation analysis (Figure 2.A). Finally, G×Y×L variance could not be assessed for all traits and remained moderate (2.7–16.4%). Generally speaking, all responses to biotic stresses were found to be influenced by G×E interactions, with the response to leaf curl being the most stable across environments.

**Figure 2.**
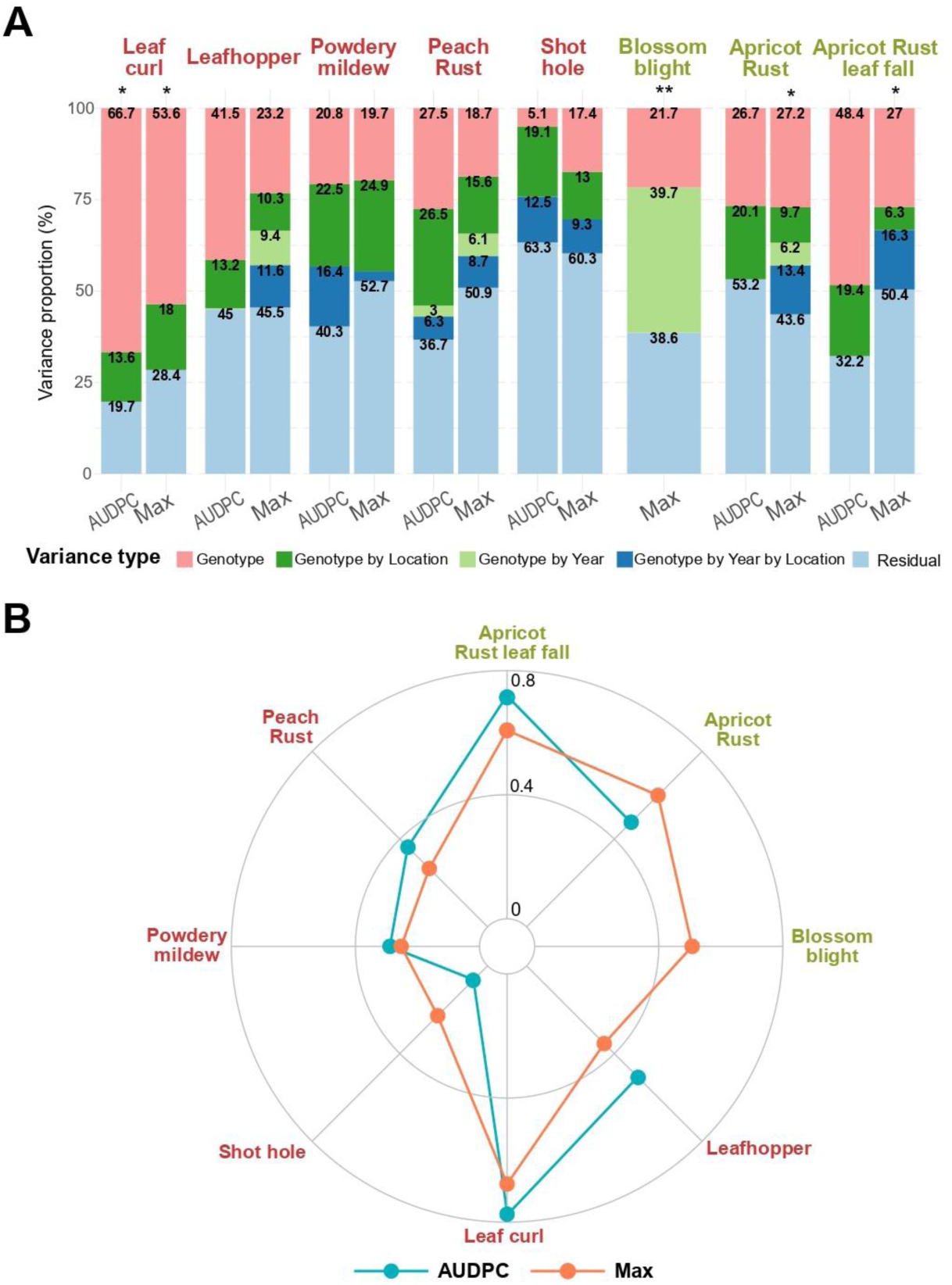
A) Stacked bar plots with the variance of the random effects of genotype, genotype by location, genotype by year and genotype by year by location interactions, and residuals; calculated from the model presenting in Eq. (2) of Supplementary Method 3 and B) broad sense heritability calculated from Eq. (3) Supplementary Method 3. *\*Biotic stress observed only one year* *\*\*Biotic stress observed only in one location*

Overall, broad-sense heritability estimates across environments on spatially corrected Max and AUDPC variables ranged from 0.07 to 0.78 for the eight pests and diseases studied (Figure 2.B). Heritability was generally higher when considering AUDPC rather than Max.

When dissecting between additive and non-additive variance, we found that the dominance variance represented up to 16.4% of the total variance, depending on the trait, the highest proportions being found for apricot rust (Supplementary Fig 5). Except for powdery mildew, the dominance variance was higher when considering Max as compared to AUDPC. A comparison of models for these across-environment variance decompositions using the BIC criterion suggested that the Additive-Dominant model was the best model choices for five traits (leafhopper Max and AUDPC, powdery mildew Max and AUDPC, and apricot rust Max).

### Genome-wide association analyses for biotic stress responses in the two core collections

We exploited genetic and G×E variance by applying a series of complementary GWAS models whereby phenotypes were considered i) across environments (G-BLUPs), ii) within environment (environment-specific BLUPs) and iii) in multiple environments. For each analysis significant, SNPs were grouped in QTLs by considering linkage blocks using the chromosome-specific LD decay values (ranging between 508 and 1620 Kbp and between 131 and 473 bp respectively in the peach and the apricot core collection, depending on the chromosome) (Supplementary Fig 6).

### Complementarity between additive and non-additive QTLs identified across environments

To exploit both additive and non-additive main genetic effects, four different models were tested on G-BLUPs, representing the different modes of action of genes: additive, dominance, recessive and overdominance (the last three have been grouped as ‘non-additive’; *see material and methods section*).

When considering the union of QTLs obtained for Max and AUDPC (*i.e.* detected for one, the other, or both variables), we identified between 10 and 131 QTLs per trait (Figure 3.A). Within the total of 341 QTLs detected, 188 were additive, 153 were non-additive, and 30 were detected with both additive and non-additive models (Figure 3.A). Except for apricot rust and rust leaf fall, more QTLs were identified with non-additive as with additive models (between 1.2 to 3.7 times more QTLs depending on the trait). Among the non-additive models, the recessive model identified the most QTLs across all traits (between 1 to 28 with an average of 8.8) (Figure 3.C and Supplementary Fig 7), while the over-dominance model identified the least (between 0 to 19 across traits, with an average of 4.9) (Figure 3.D and Supplementary Fig 7). However, it should be reminded that dominant and recessive models are only differentiated regarding the effects of the reference allele. Therefore, QTLs detected with both types of models can correspond to a recessive or dominant mode of action.

**Figure 3.**
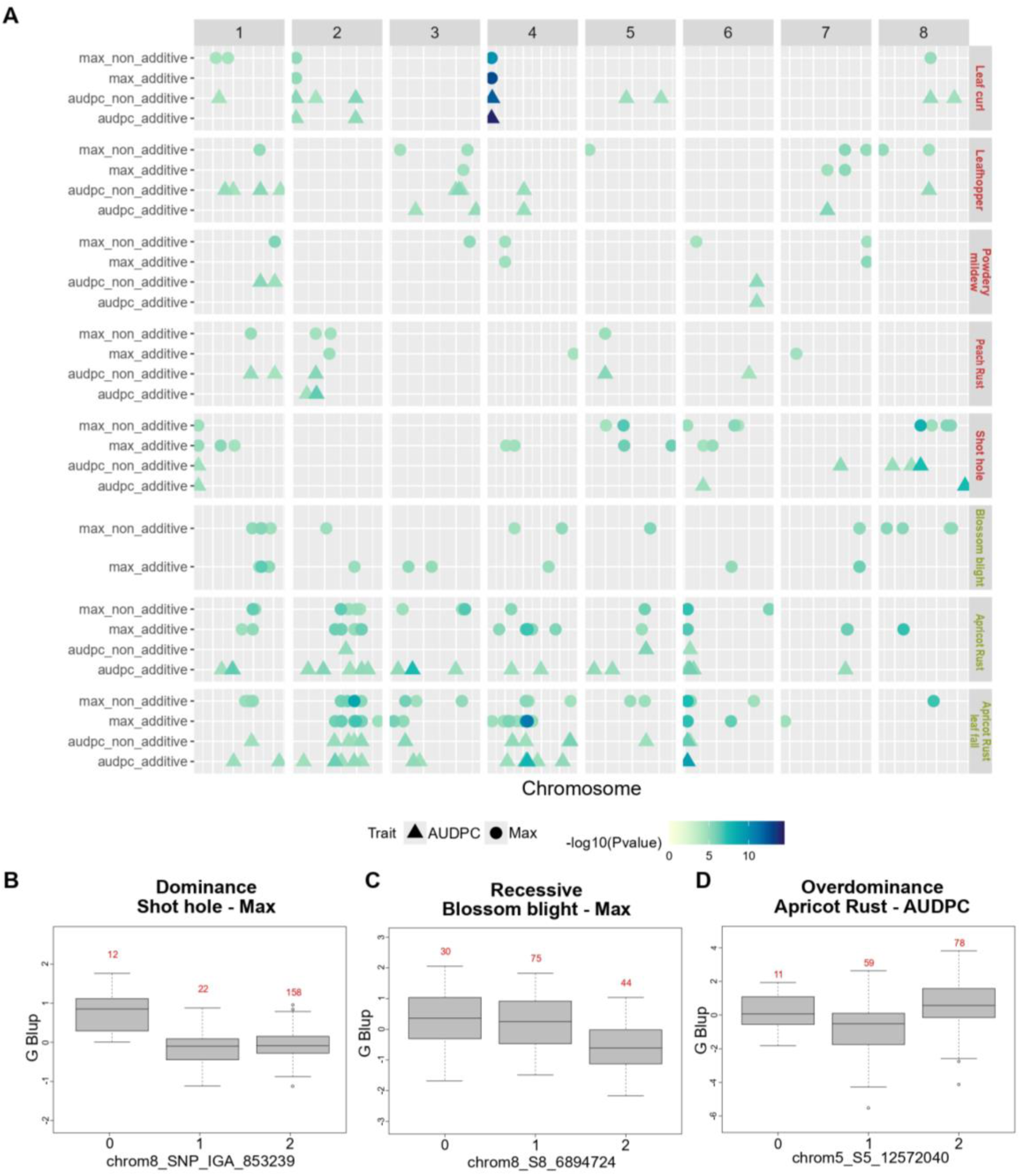
GWAS results on main additive and non-additive genetic effect with A) QTL summary showing the physical position of GWAS significant SNPs. Each horizontal line contains QTLs of one biotic stress, organized by additive or non-additive analysis and either for the maximum of damage score (called Max) or the AUDPC. The dot colours are proportional to the -log_10_(*P*-value) of the QTL top SNP. The x-axis indicates the physical positions on the peach or apricot genome. Boxplots illustrating the effect of a (B) dominant, (C) recessive and (D) overdominant SNP with a significant association for respectively shot hole (Max), blossom blight (Max) and apricot rust (AUDPC).

We found respectively 271 and 119 QTLs when considering Max and AUDPC variables with a total of 41 overlapping QTLs. Apricot rust was the disease for which Max and AUDPC QTLs overlapped the least (with less than 1% of overlapping QTLs) whereas leaf curl was the disease with the most overlap (with 26% of overlapping QTLs) (Figure 3.A).

Overall, the non-additive GWAS analysis allowed identifying complementary QTLs that were not detected by the additive-GWAS analysis, with 96 and 45 QTLs being exclusive to non-additive models respectively for Max and AUDPC.

### Identification of environment specific QTLs with single-environment GWAS

In order to study environment-specific effects, we implemented single-environment GWAS using single-Locus Mixed-Models on environment-specific BLUPs. We identified between 33 and 568 significant SNPs depending on the trait, corresponding to a total of 801 QTLs (union of results on Max and AUDPC variables) (Figure 4).

**Figure 4.**
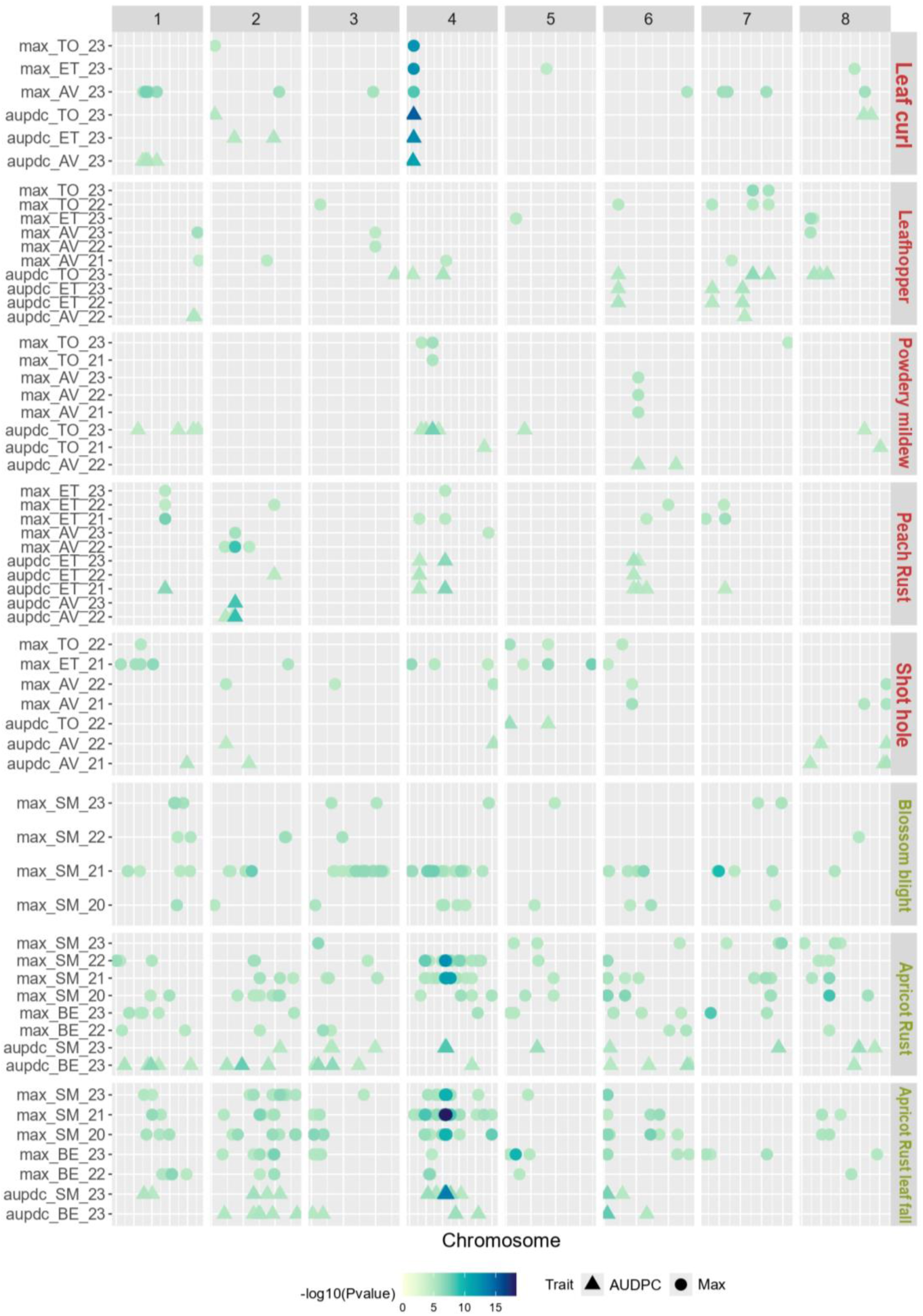
QTL summary showing the physical positions of GWAS significant SNPs for environment-specific models. Each horizontal line contains QTLs of one biotic stress, organized by environment and either for the maximum of damage score (called Max) or the AUDPC (except for blossom blight, where AUDPC was not available). The coloured dots are proportional to the -log_10_(*P*-value) of the QTL top SNP. The x-axis indicates the physical positions on the genome (peach or apricot depending on trait).

The identity and number of associations were greatly dependent on the target environment (62.5% to 100% environment-specific QTLs depending on trait) (Figure 4). Leaf curl was the only trait for which a QTL was identified in all investigated environments and for both AUDPC and Max variables. Apart from this QTL, located on chromosome 4, the other 25 QTLs associated to leaf curl were detected in only one environment. For apricot rust and rust leaf fall, some QTLs have also been identified in all the environments studied when considering AUDPC (representing respectively 5 and 3% of all the QTLs identified). With a total of 6 QTLs detected in 2 different environments and 3 QTLs detected in 3 different environments (total of 5 environments considered), peach rust was the trait with the highest proportion of stable QTLs. On the opposite, blossom blight was the disease with less stable QTLs as all of them were found to be environment-specific. For all traits, QTLs associated to Max appeared to be more stable than those found with AUDPC (22% and 0.09% of QTLs detected in more than one environment, respectively). Finally, among the QTLs identified, between 2 and 46 QTLs overlapped between Max and AUDPC depending on the biotic stress.

### Modelling QTLs with G×E interaction effects using MTMM and Meta-GWAS

To statistically distinguish between stable QTLs across environments and those underlying G×E interactions, we performed i) a multi-environment analysis on environment-specific BLUPs using the fully parameterized multi-trait mixed model (MTMM)[29] comprising three sub-models (‘common model’, ‘gei model’ and ‘full model’) (Supplementary Method 6), and ii) a Meta-GWAS with the MetaGE model [34](Supplementary Method 7). MTMM disentangle common effect markers from interactive markers and MetaGE jointly analyses single-environment GWAS results to identify QTLs with minor effects which might go undetected in single environment analyses, as well as markers with significant effects across multiple environments.

MTMM GWAS and Meta-GWAS analyses led to the identification of respectively 209 and 226 QTLs significantly associated when considering all traits (union of Max. and AUDPC), with 1 to 61 QTLs per trait identified with MTMM and 1 to 54 QTLs per trait identified with Meta-GWAS (Figure 5). From those, 107 QTLs overlapped between MTMM and MetaGE approaches.

**Figure 5.**
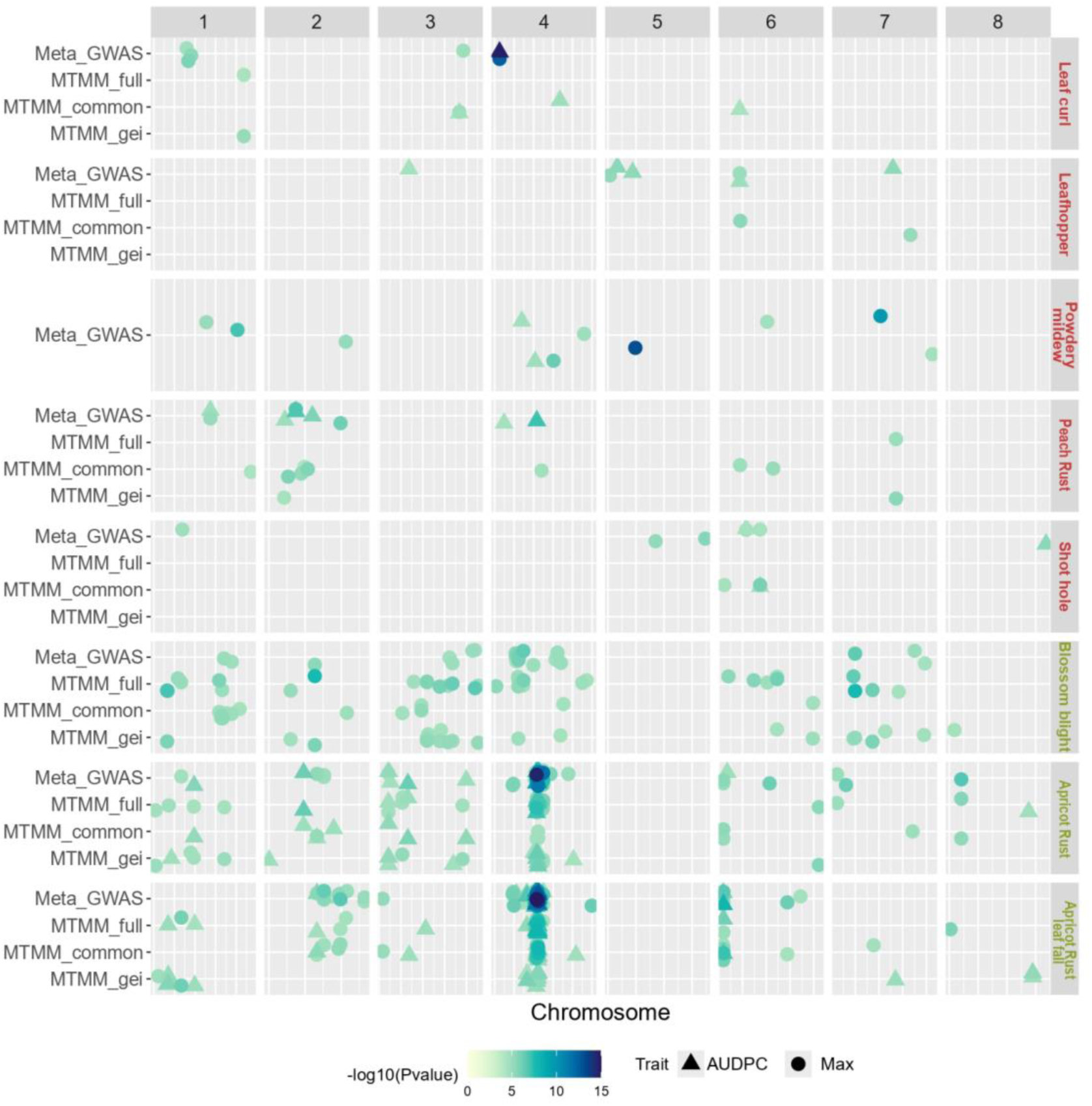
QTL summary showing the physical position of GWAS significant SNPs for the two models dissecting the G×E interactions (MTMM and Meta-GWAS). Each horizontal line contains QTLs of one biotic stress, organized by GWAS model (Meta-GWAS and the 3 sub-model of MTMM: full, common and gei). The dot colours are proportional to the -log_10_(*P*-value) of the QTL top SNP and the dot shape represent the considered variable (Max or AUDPC). The x-axis indicates the physical positions on the peach or apricot genome.

MTMM analysis could not be performed on 4 variables (AUDPC of peach rust and leafhopper, and on both Max and AUDPC for powdery mildew). For the remaining, we identified 75 and 67 QTLs respectively for ‘common’ and ‘gei’ models (union of Max. and AUDPC, Figure 5). For most traits, more QTLs have been identified with the ‘common’ model than with the two other MTMM models (between 50 to 100% of QTLs detected). However, for blossom blight (Max), apricot rust (AUDPC and Max) and apricot rust leaf fall (Max) 53 to 65% of QTLs were identified with the ‘gei’ model. For these ‘gei’, interactive QTLs, allelic effects of the top SNP ranged from negative to highly positive depending on environment.

When performing Meta-GWAS, we found a large amplitude of z-scores - a summary statistics of the marker’s *P*-value of the sign of the marker’s effects - for most SNPs. We observed only rare sign inversions, indicative of SNPs with the same qualitative effects (*i.e.* same direction), but quantitatively different effects across environments (Supplementary Fig 8). A remarkable exception was found for blossom blight, with 61% of the significant SNPs presenting a z-score sign inversion between the 4 environments considered (Supplementary Fig 8.F). Interestingly, we detected 29 QTLs with Meta-GWAS that had not been detected with single environment GWAS (Figure 4 and Figure 5).

### Identification of high confidence QTLs and quantification of their contribution to G and G×E interactions

When considering all traits and models tested, 101 QTLs were detected for peach and 606 for apricot. To retain the most relevant QTLs, high confidence QTLs were defined as those detected in at least 50% of the additive GWAS models tested. We retained up to 32 high confidence QTLs depending on the trait (Supplementary Table 2). Blossom blight was the only disease for which no confidence QTLs could be identified. High confidence QTLs overlapping between AUDPC and Max have been found for leaf curl (1 QTL located on chromosome 4), apricot rust (1 QTL located on chromosome 4) and apricot rust leaf fall (18 QTLs located on chromosome 2, 4 and 6) (Supplementary Table 2).

For each trait, we calculated the proportions of G, G×L and G×Y variance captured by the high confidence QTLs (considered jointly when several confident QTLs were identified for a given pest or disease). The high confidence QTLs captured between 9.6 and 64.6% of the additive genetic variance depending on the biotic stress (Supplementary Fig 9). These QTLs also explained variable portions of G×L interactions (up to 54.4%) and to a lesser extent, of G×Y interactions (up to 29.3%) (Supplementary Fig 9). Interestingly, in apricot rust high confidence QTLs explained a higher proportion of the G×L variance than of the genetic variance (Max variable) (Supplementary Fig 9).

### Local LD and annotation analyses within high confidence QTLs towards identifying candidate genes

Based on the association results, we searched within high confidence QTLs for candidate genes with predicted functions linked to host defence or response mechanisms. As a preliminary step, we readjusted peach QTL intervals to account for the strong LD observed in peach by considering the local LD between the top SNP of each high confidence QTL and all SNPs belonging to this QTL. We considered that a SNP was independent from the top SNP when is r^2^ to top SNP was below 0.2 (Figure 6.A and B for Leaf_curl_4 and Supplementary Fig 10 for the other peach QTLs). This enabled us to narrow down the list of candidate genes for peach, with an average of 34.6% of genes with a relevant function located in the readjusted QTL intervals (Table 1 and Supplementary Table 3).

**Figure 6.**
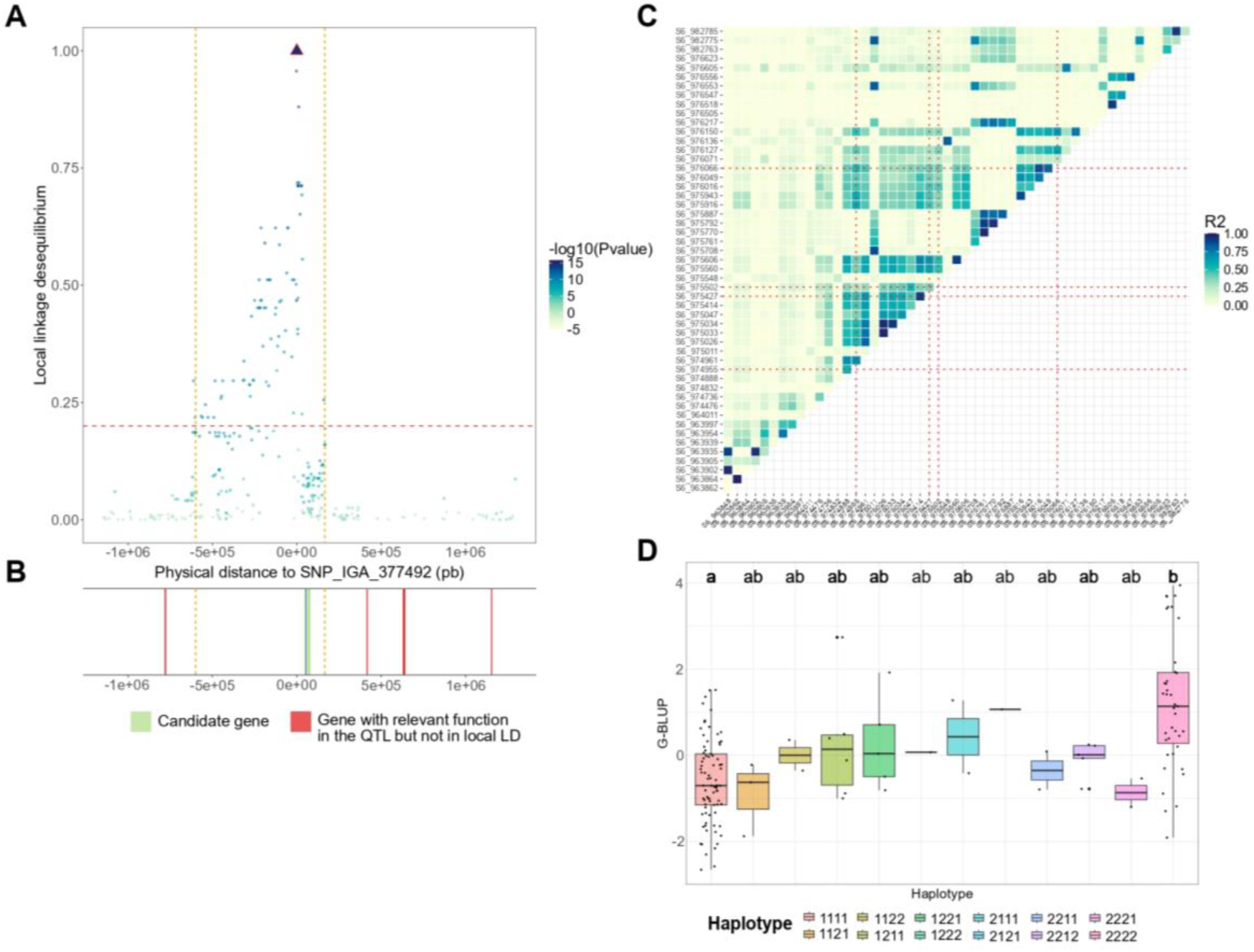
Fine characterization of two high confidence QTLs. A) Representation of the local LD between the top SNP of QTL ‘Leaf_curl_4’ and all other SNPs of the QTL. The X-axis represents the physical distance from the top SNP, while the Y-axis represents the r^2^, corrected by the relatedness and structure, between SNPs and the top SNP. Each dot represents a SNP which is coloured according the p-value of Meta-GWAS. The vertical orange dot lines represent the QTL interval readjusted using the local LD. B) genes with relevant functions located in the QTL, with candidate genes in green and genes located in the QTL but not in local LD with the top SNP in red. B) Relevant genes located in the QTL, with candidate genes in green and relevant genes located in the QTL but not in local LD with the top SNP in red. C) Heatmaps of pairwise LD estimates within the genomic window around candidate gene ‘PruarM.6G017900’. Top SNPs of the different high confidence QTLs are shown with red dot line and D) G-BLUPs distributions of apricot rust Max according to the haplotype constituted by the SNPs ‘S6_974955’,’S6_975427’,’S6_975502’ and ‘S6_976066’. Statistical differences between groups were computed by pairwise Wilcoxon tests.

**Table 1.**
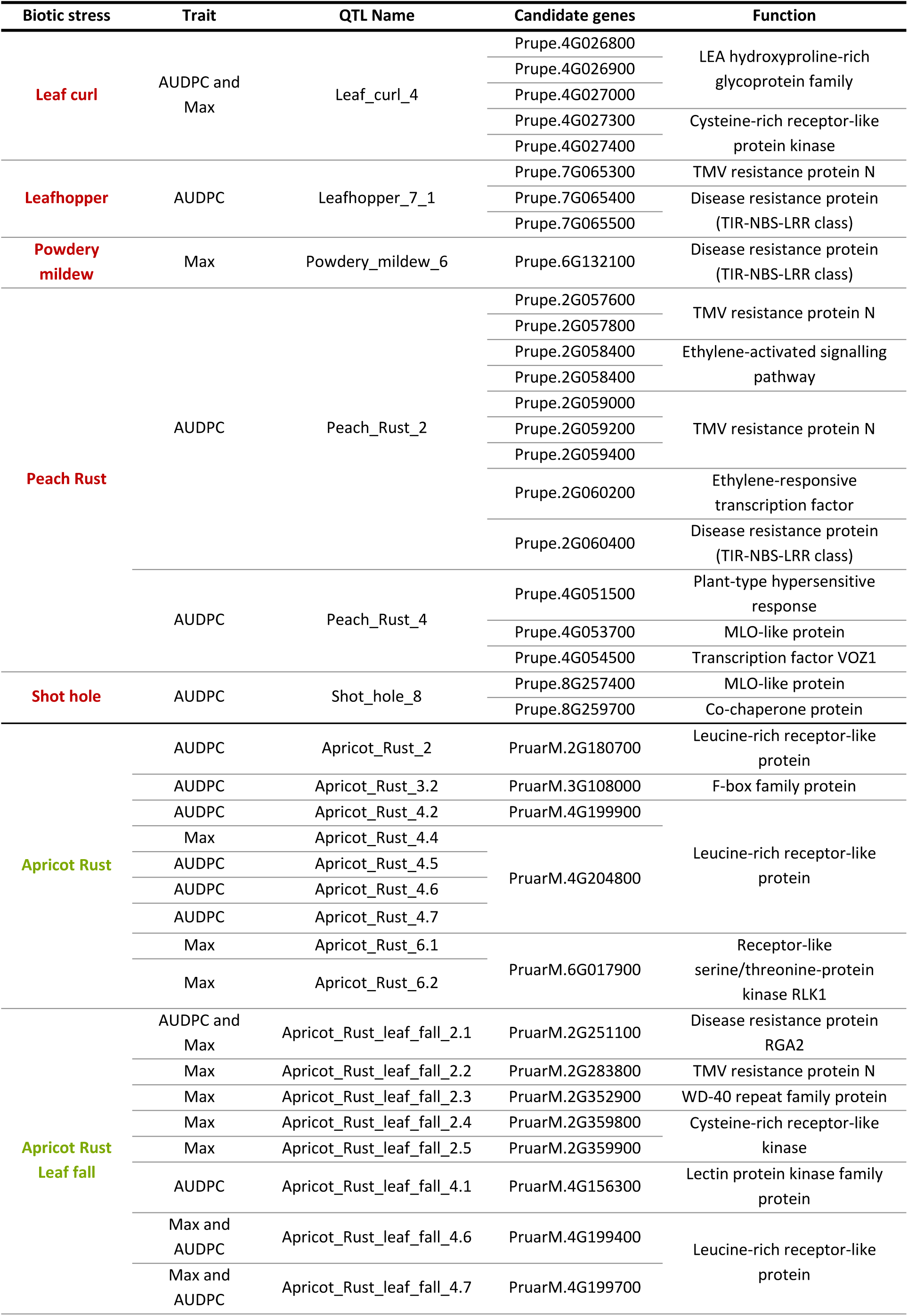
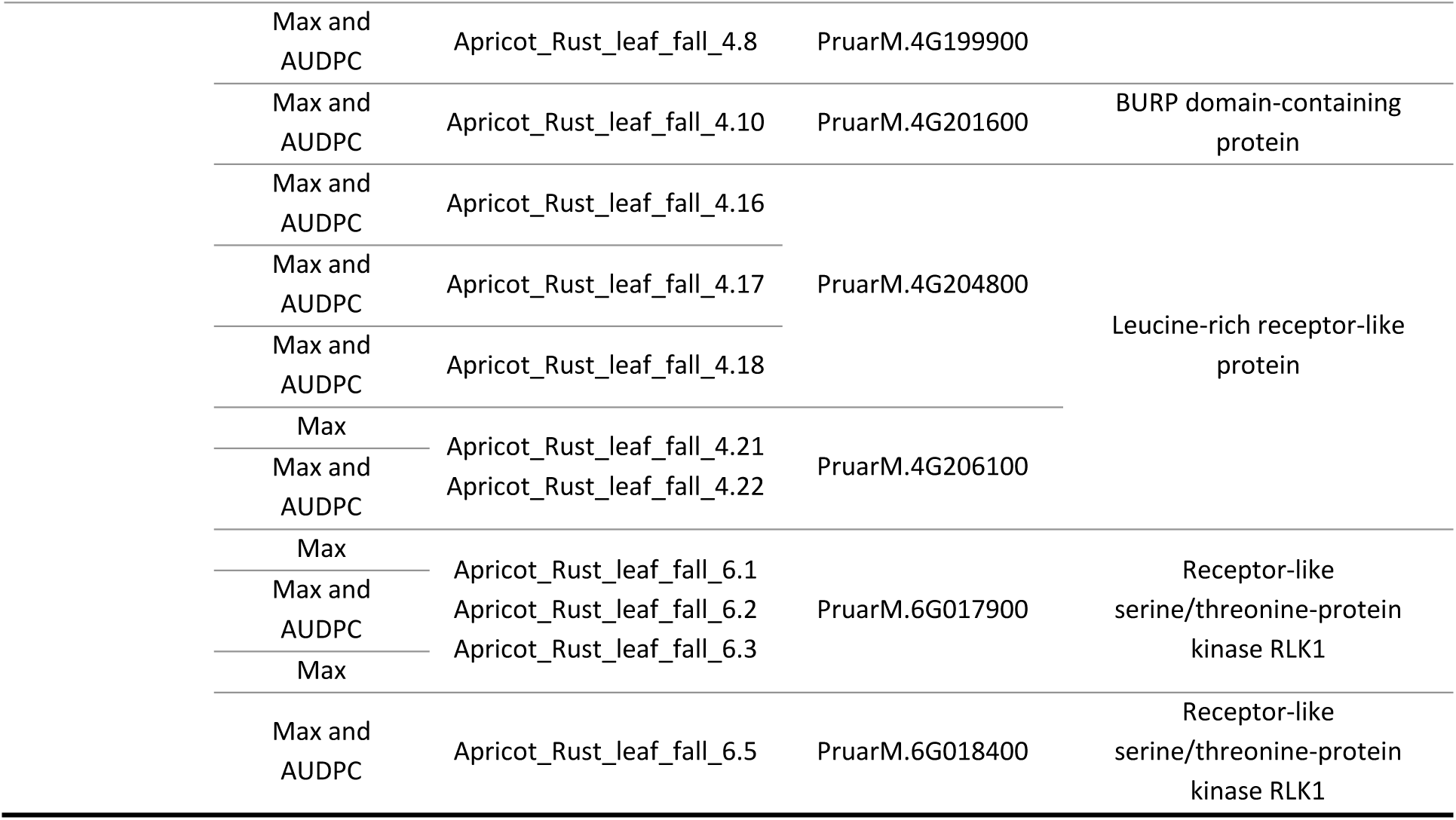
List of candidate genes for the high confidence QTLs

A total of 87 candidate genes have been identified across the 60 high confidence QTLs, with up to 9 candidate genes for the peach rust QTL on chromosome 2 (Table 1). Among the candidate genes, seven were linked to TMV resistance protein, seven to Leucine-rich receptor-like protein kinase family protein, four to cysteine-rich receptor-like protein kinase, four to TIR-NBS-LRR disease resistance protein and two to MLO-like protein (Table 1). For some QTLs associated to apricot leaf rust, leafhopper and shot hole no candidate genes could be clearly identified.

Interestingly, three apricot candidate genes, ‘PruarM.6G017900’, ‘PruarM.4G204800’ and ‘PruarM.4G206100’, were identified as covering several QTLs (Table 1). We calculated the local LD in these three zones (Figure 6.C. and D, Supplementary Fig 11) and observed two different patterns: for ‘PruarM.6G017900’ (Figure 6.C) and ‘PruarM.4G204800’ (Supplementary Fig 11.A) a relatively high LD block overlapped precisely with the underlying candidate gene. These results suggested that the QTLs identified in these different genes were not independent. In contrast, ‘PruarM.4G206100’ appeared to be located in a moderate LD region with no clear LD blocks (Supplementary Fig 11.B). Max G-BLUPs distributions according to the haplotype constituted by the top SNPs identified in these three genes were plotted (Figure 6.D, for top SNPs belonging to ‘PruarM.6G017900’ and Supplementary Fig 11.C and D for ‘PruarM.4G204800’ and ‘‘PruarM.4G206100’). For both genes ‘PruarM.6G017900’ and ‘PruarM.4G204800’, the two most frequent haplotypes, had significantly different G-BLUPs values (Wilcoxon test *P*-value < 0.05) (Figure 6.D and Supplementary Fig 11.B).

## Discussion

The main objective of this study was to dissect the genetic architecture of a series of traits reflecting the response to natural infections in multiple environments in peach and apricot. We specifically aimed at deciphering G×E interactions, identifying candidate genomic regions, and comparing the results obtained in these two species.

### Biotic stress response is highly influenced by G×E interactions

Our results revealed that the expression of pest and disease susceptibility was under a strong influence of the environment *per se* but also of G×E interactions (Figure 1, Figure 2, Supplementary Fig 1, Supplementary Fig 2 and Supplementary Fig 3). Variance decompositions (Figure 2.A) and the pairwise genetic correlations (Supplementary Table 1) indicated the prevalence of the G×E interactions over G effects for several traits. Similar patterns have been found in stone fruit diseases, such as in peach brown rot, where skin tolerance to the fungal infection appears to be under high seasonal influence [50,52]. In a study on apricot bacterial canker, Omrani *et al.* (2019) pointed to the relevance of controlling G×E interactions for accurately estimating genetic effects in the face of environmental variations in multi-year data [49]. In our study, G×E interactions were generally driven more by location than by year effects, which was supported by phenotypic and GWAS analyses. With single-environment GWAS, we identified a total of 567 environment-specific QTLs and 135 environment-shared QTLs with a higher overlap between QTLs identified from each location than from QTLs identified from a given year. Interestingly, the QTLs identified with the multi-environment models (MTMM and Meta-GWAS) were mostly interactive which changes in direction (‘antagonist’) or intensity (‘differential’) according to the environment. Our results thus indicate that data on specific climatic conditions have the potential to reveal additional regions linked to biotic stress response. We thus recommend evaluating the same panel - or highly overlapping panels including several common genotypes - in multiple geographic areas with contrasted environmental conditions.

It should be reminded that the functional validations of G×E interaction QTLs are still rare and requires challenging tests such as gene cloning [61]. Finally, a better characterization of the environments through envirotyping could allow to identify which environmental variables modulate the QTLs effects and to group locations accordingly for common adapted varieties [27].

### A focus on peach leaf curl and apricot blossom blight

Among the seven biotic stresses studied, leaf curl in peach and blossom blight in apricot stood out from the others by presenting two distinct typologies: the former being little influenced by environments and G×E interactions and the latter, by contrast, being strongly influenced by environmental effects. Our study provides the first GWAS analysis on these two major stone fruit diseases.

Apricot blossom blight variation was found to be controlled by a high number of environment-dependent QTLs, indicating a complex genetic determinism. Variance decompositions and GWAS analyses pointed towards predominant G×Y effects on this disease. In fact, all QTLs identified with single-environment models were year-specific and respectively 65% and 61% of the QTLs identified with MTMM and meta-GWAS presented allelic effects modulated by year (*i.e.* interactive QTLs). Due to these important fluctuations between years, no high-confidence QTLs could be identified with our method. It has been shown that the development of Monilinia on apricot is tightly linked to the interaction between phenology and meteorological conditions, where temperature and humidity during blossom play a major role [62,63], and knowing that the petals being the entry point for the fungus [64,65]. Our dataset was filtered prior to analyses to avoid strong pheno-climatic bias by taking out blossom blight incidence scores for trees having insufficient flower density and Monilinia climatic index in a given year [66]. Despite decent heritability values obtained for apricot blossom blight (0.51; Figure 2.B), we might need to better account for the epidemiological context to further study the genetic basis of that disease.

In contrast to apricot blossom blight, leaf curl damage was a highly heritable trait underlined by low environmental and G×E effects (Figure 2 and Supplementary Table 1). This low environmental impact could be partly explained by a homogeneous leaf curl pressure across the targeted geographic areas, but also within fields and along the susceptibility period (leaf emergence stage). On the opposite of blossom blight, where the susceptible stage (*i.e.* flowering) lasts for a few days to a week, the infection period for leaf curl lasts over several weeks [67] meaning that phenology-driven avoidance is rare. In line with the phenotypic analyses, we found a highly stable QTL located on chromosome 4 (identified with all GWAS models tested apart from MTMM). 23 further QTLs have been identified for leaf curl, with an intermediate level of confidence, and mostly driven by location-specific effects (19 location-specific QTLs identified with single-environment models). These results suggest that low susceptibility to leaf curl is governed by a major QTL combined with several minor genes having weaker and environment-specific effects.

In terms of experimental design, we would recommend studying blossom blight incidence over several years in a location with overall high incidence (*e.g.* upper Rhone Valley in France), to capture for each accession a season during which the tree was actually confronted to the risk. Regarding leaf curl, several locations might be necessary only to capture the minor-effect genes that act as a complement to the major QTL. In terms of breeding strategies, MAS seems to be particularly well adapted to leaf curl, whereas for blossom blight, combining improved epidemiological risk prediction with genomic selection would be a promising strategy.

The rest of the biotic stresses appeared to be more nuanced with no strong typology emerging. We highlighted a location effect but the relative classification of accessions is relatively well preserved between sites (*i.e.* rank inversions between accessions are not so common). On average, 23% of the QTLs identified with single-environment GWAS were ‘environment-shared’. Finally, the majority of ‘interactive’ QTLs identified presented a ’differential’ profile which suggest that some QTLs have effects in some environments and not in others, but true inversions of QTL effects are less common.

### Usefulness of AUDPC and Max variables to study biotic stress response: identification of complementary regions

We decided to compare Max and AUDPC, two key metrics to capture the complexity of disease dynamics in naturally infected fruit tree orchards. The maximum of damage score provides a clear indicator of ‘absolute’ vulnerability to biotic stresses but overlooks disease progression over time in different epidemiological contexts.

Although we found high correlations between Max and AUDPC for all biotic stresses studied, with broadly similar heritability (Figure 1, Figure 2.B and Supplementary Fig 2), we identified between 31 to 97 new QTLs with AUDPC depending on the GWAS models. Importantly, among the high confidence QTLs, 32% where only detected using AUDPC. Overall, QTLs detected with Max were more stable while those identified with AUDPC were more location-or year-specific. This underlines the complementary nature of the two metrics. Although widely used in other crops, mostly under controlled infection conditions, the use of AUDPC has increased in recent years for assessing quantitative disease resistance in fruit trees and for evaluating disease management practices [68–70]. Some studies have even highlighted the dynamics of genetic effects during pathogen colonization with a distinction between early and late response and temporal pattern in effects of QTLs [69].

Because AUDPC requires a large effort to provide repeated measures at multiple time points of the season, we recommend implementing this metric for diseases for which the onset and duration of infection is critical for production, such as apricot rust, or for polycyclic disease [71]. Indeed, early summer defoliation caused by rust has a strong impact on flowering capacity next year. We showed that among the high confidence QTLs identified for apricot rust and rust leaf fall, 25% were AUPDC specific and only 37% overlapped between AUDPC and Max. Therefore, AUDPC enables to better distinguish accessions with a high and continuous level of infection at a critical time point of the season, which is not possible with Max.

### High confidence QTLs reveal the role of basal and host-specific resistance shaping the response to biotic stress in stone fruit trees

Among the QTLs detected with single environment models, respectively 79%, 77% and 54% were also detected with GWAS model focusing on the main genetic effect, by the multi-environment GWAS (MTMM) and with meta-GWAS. In total, these different GWAS models enabled the detection of respectively 38, 79 and 29 QTLs that had not been detected with single environment GWAS. The findings in our study support combining single-and multi-environment models to improve the statistical power, the reliability and robustness of GWAS. Similar results have been obtained in the case of QTL detection for drought response in maize where the incorporation of the interaction with the environment improved the power of GWAS and enabled to find QTL that are significant in a broad range of environments [72]. By conducting GWAS with multiple methods, we narrowed down the list of relevant regions to a total of 60 ‘high confidence QTLs’ across traits. These QTLs explaining in average 36.8% of the genetic variance, and for apricot rust they also explained 53.6% of the genotype by location interaction variance.

To our knowledge, for most of the pests or diseases studied here, resistance or tolerance mechanisms have never been investigated by GWAS analysis neither by QTL mapping, as is the case for leafhopper, shot hole, blossom blight, peach and apricot rust. Previous analyses on peach powdery mildew have already identified QTLs and candidate genes (Vr2 and Vr3) located on chromosomes 8 and 2 [73–75], but none of them overlap with the high confidence QTL identified here. The high confidence QTL we identified for leaf curl on LG4 was not reported previously. A unique linkage analysis study on leaf curl in an interspecific F1 progeny between a peach cultivar and *P. davidiana*, a peach related species, reported two consistent QTLs located on linkage groups 3 and 6 [76]. QTLs were also identified in these regions by MTMM and Meta-GWAS but they were not categorized as ‘high confidence’. In contrast to a QTL-mapping population where the genetic background is homogenous, major alleles might go undetected due to low frequency in a core collection. This may explain the low overlap between our results and those already reported. The type and number of markers included, the approaches used for detecting QTLs or the difference in phenotyping techniques might also contribute to some discrepancies between studies.

Interestingly, the majority of candidate genes identified belong to the Leucine-rich repeat (LRR)-containing receptors (LRR-CRs) family gene which represents the largest class of plant resistance genes (R genes) [77,78]. LRR-CRs can be classified in three main sub-families: LRR receptor-like kinase (LRR-RLK), LRR receptor-like protein (LRR-RLP) and nucleotide-binding-site LRR (NBS-LRR) [79,80]. In our study we identified candidate genes belonging to all the categories with genes related to i) LRR-RLK identified for apricot rust and rust leaf fall (PruarM.6G017900 and PruarM.6G018400), ii) LRR-RLP identified for apricot rust and rust leaf (PruarM.2G180700, PruarM.4G204800, PruarM.4G199400, PruarM.4G199700, PruarM.4G199900, PruarM.4G204800 and PruarM.4G206100) and iii) NBS-LRR identified, for leafhopper (Prupe.7G065400 and Prupe.7G065500), powdery mildew (Prupe.6G132100), peach rust (Prupe.2G060400). RLKs and RLPs have been identified to function as cell surface pattern-recognition receptors (PRRs) which detect pathogens-associated molecular patterns (PAMPs) and lead to the activation of the pattern-triggered immunity (PTI), the basal defence responses to pathogens [79,81]. We also identified a high number of genes annotated as ‘TMV resistance protein N’ (Prupe.7G065300 identified for leafhopper, Prupe.2G057600, Prupe.2G057800, Prupe.2G059000, Prupe.2G059200 and Prupe.2G059400 identified for peach rust, and PruarM.2G283800 for apricot rust leaf fall). The tobacco N gene, conferring resistance to tobacco mosaic virus (TMV) has been identified as TIR-NBS-LRR protein type [82]. It has been shown that NBS-LRR proteins conferred resistance to diverse pathogens, including fungi, oomycetes, bacteria, viruses and insects. By recognizing pathogen effector molecules, NBS-LRR proteins lead to the activation of defence responses and play a fundamental role in the effector-triggered immunity (ETI), the second pathogen-sensing mechanism in plants [81]. ETI usually induces programmed cell death at the infection site, known as the hypersensitive response (HR), and thus locally limits pathogen spread. NBS-LRR proteins are highly variable and subject to balancing selection, which could contribute to the high specificity of these effectors [79]. Thus, our results point towards a prevalent role of host specific (narrow spectrum) and non-specific (large spectrum) resistance mechanisms among the diseases studied here.

### Similar, yet divergent: difference in the architecture of biotic stress response between peach and apricot in the light of their biological features

Peach and apricot are closely related species belonging to the *Rosaceae* family and the genus *Prunus*. They present different reproductive characteristics and demographic history [83]. Endowed with a self-incompatibly system, apricot is a cross-pollinating species [84], whereas peach is a preferentially self-pollinating species [85]. This results for apricot in a high number of effective recombination events and a rapid expected LD decay over the genome, whereas the opposite pattern is expected for peach [86]. In our study, global mean LD blocs over all chromosomes are about 3300 times longer for the peach core collection compared to the apricot core collection. In fact, previous association studies have already highlighted a LD decay over very short distances (100 to 200 bp) in apricot (based on a population comprising an important part of the material used in presented work) [48,49]. In a similar way for peach, other studies have found a significant LD decay at 5 Kbp to 1.8 Mbp (depending on the population), confirming high levels of LD over long distance for that crop [44,87]. QTL intervals were therefore particularly small for apricot (max 946 Bp), with most of them being defined by the top SNP. Minding a sufficient number of markers, a higher mapping resolution makes association analyses more powerful to target causal polymorphisms in apricot, facilitating the identification of promising candidate genes [88]. For rust tolerance, we identified three genes covering several confidence QTLs in apricot, along with favourable haplotypes (Figure 6.D and Supplementary Fig 11). Results on apricot strongly contrast with those obtained in peach, where QTLs were up to max 3240 Kbp in size, and contained 52 to 528 genes. Using local LD values allowed to refine QTL sizes and to narrow down the list to 23 candidate genes.

In our study, rust was the only disease able to affect both peach and apricot. Rust diseases have a fungal origin and affect many crops, and typically present a high specificity to their host [89]. *Tranzchelia discolor* (Fuckel) has been shown to attack peach, almond, plum, apricot, and cherry [90,91]. For both species, a high confidence QTL was identified on chromosome 2.

Finally, we identified a total of 123 new QTLs using non-additive models, indicating that non-additive effects play a significant role in biotic stress response for both species. In proportion, more non-additive QTLs were found in peach (56%) than in apricot (39%). Although peach is a self-compatible species with a low level of heterozygosity, this result indicates that both additive and non-additive genetic effects are important in the genetic architecture of the traits under study, as already been shown for leaf curl susceptibility inheritance [92]. In almond, another cross-pollinating *Prunus* species, non-additive GWAS conducted on agronomic traits led to the identification of 13 QTLs with only one having additive effect [83]. Harnessing non-additive variance might thus be valuable to better exploit genetic variance, which is particularly relevant in clonally propagated crops.

### Conclusion and prospect

This study provides a thorough characterization of the genetic variation underlying resistance and tolerance to several pests or diseases in peach and apricot via observations over multiple environments under natural conditions. By quantifying and exploiting G×E interactions, we found a majority of environment-specific or interactive QTLs. For future experiments, we recommend considering the combination of pathogens which are specifically expressed in each target location to optimize the choice of experimental sites.

Our results emphasize that, while accounting for pedo-climatic adaptation and market requirements, future breeding should focus on adaptation to the biotic profile of the target production area. Developing new peach and apricot cultivars is a time-consuming process because of the long juvenile stage, large plant size, and the multi-year recording of phenotypic performance [93]. As a next step, the effects of the allelic variants identified in this study need to be validated before incorporating them in MAS strategy to efficiently screen breeding materials and optimize the multi-trait and multi-environment selection schemes. Another promising strategy to overcome the limiting aspects of GWAS would be the use of genomic prediction models for a better exploitation of weak effects underlying quantitative resistances and tolerances [94,95].

## Material and methods

### Plant material and trial description

The plant material and experimental design used in this study were previously described in [96]. To sum up, we used two core collections of respectively 206 peach (*P. persica and 3 relatives: P. davidiana, P. mira,* and *P. kansuensis*) and 150 apricot (*P. armeniaca*) accessions coming from multiple geographical origins with significant number of accessions of historical and regional importance, landraces and elite materials. The peach core collection also includes ornamental cultivars, few accessions from close-related species and interspecific crosses.

All orchards are located in South-East France, in three locations for peach: Étoile-sur-Rhône (44.8°N/ 4.9°E, Drôme), Avignon (43.9°N/4.8°E, Vaucluse) and Torreilles (42.8°N/3.0°E, Pyrénées-Orientales), respectively called ET, AV and TO; and in two different locations for apricot: Saint-Marcel-lès-Valence (45.0°N/4.9°E, Drôme) and Bellegarde (43.7°N/4.5°E, Gard), respectively called SM and BE. Climate ranges from Mediterranean to semi-continental. Peach and apricot orchards are composed of two and five randomized complete blocks respectively, each block containing one replicate per genotype. All accessions have been grafted on ‘Montclar® Chanturgue’ peach rootstocks. First leaf emergence was in 2019 for peach and in 2018 for apricot.

The orchards were managed under integrated plant protection during the first two years after the plantation with a progressive reduction of phytosanitary protection thereafter. Overall, only punctual treatments were applied if the survival of the orchard was at stake for peach orchards since 2023, and except for the protection against the psyllid vector of the Apricot Chlorotic Leaf Roll (ACLR) in Bellegarde, no protection was applied for apricot orchards since 2020.

### Phenotypic data collection

We monitored the damages caused by four diseases and one pest on peach in 2021, 2022 and 2023 and by two diseases on apricot in 2020, 2021, 2022 and 2023 (Table 2). For the rest of the analysis, the term of environment refers to the combination of a location and a year. All field data were collected using the ‘Field Book’ Android application [97].

**Table 2.** Summary of pests and diseases monitored in this study

|  | Pest and Disease | Organism | Pathogen | Organ | Number of environments observed | Number of environments considered |
| --- | --- | --- | --- | --- | --- | --- |
| Peach | Leaf curl | Fungi | *Taphrina deformans* | Leaves | 3 (3 locations and 1 year) | 3 (3 locations and 1 year) |
|  | Leafhopper | Insect | *Asymmetrasca decedens* | Leaves | 6 (3 locations and 3 years) | 6 (3 locations and 3 years) |
|  | Powdery mildew | Fungi | *Podosphaera pannosa* | Leaves | 8 (3 locations and 3 years) | 5 (2 locations and 3 years) |
|  | Rust | Fungi | *Tranzschelia* spp. | Leaves | 6 (2 locations and 3 years) | 5 (2 locations and 3 years) |
|  | Shot hole | Fungi | *Wilsonomyces carpophilus* | Leaves | 9 (3 locations and 3 years) | 4 (3 locations and 2 years) |
| Apricot | Blossom and twig blight | Fungi | *Monilinia* spp. (mostly *Monilia laxa*) | Flowers and twigs | 4 (1 location and 4 years) | 4 (1 location and 4 years) |
|  | Rust | Fungi | *Tranzschelia* spp. | Leaves | 6 (2 locations and 4 years) | 6 (2 locations and 4 years) |

The assessment of the damages caused by all these pests and diseases was based on the visual estimation of symptom incidence (percentage of damaged leaves). Different rating scales were used to best capture the phenotypic expression of each pest or disease (Supplementary Method 1).

Except for blossom blight which was assessed around 30 days after blooming date and for which the shoots infected by *Monilinia* spp. were removed, repeated scoring was undertaken for the different pests and diseases to monitor the evolution of contamination throughout the seasons in the different orchards.

### Genotyping and resequencing

Genotyping of the peach core collection was performed on 192 individuals using the higher density IRSC 16K SNP array (https://www.rosaceae.org/Analysis/431). The genotyping dataset was filtered to retain SNPs with call rate per marker > 90%, missingness per individual < 50%, heterozygosity ≤ 98% and MAF >5% which resulted in a final set 13,680 markers.

The apricot core collection was sequenced with the Illumina HiSeq 2000 NGS technique. The sequence alignment was performed on the third version of the ‘Marouch’ genome [41]. The dataset was first filtered with a missingness per individual < 50% and only biallelic SNPs with more than 10 reads deep/SNP have been conserved. Finally, the dataset was filtered to retain SNPs with call rate per marker > 95%, heterozygosity ≤ 98%, MAF >5% and a removal of duplicated SNPs. This dataset consisted in 207,000 markers.

For both datasets, the imputation of missing data was done using the most frequent allele.

For the datasets used in non-additive GWAS, we included two more filtering steps to retain SNPs with three genotypic classes and with a minimum genotypic class frequency higher than 5%. These datasets consisted in respectively 10,082 markers for peach and 25,508 markers for apricot.

### Statistical analysis of phenotypic data

All statistical analyses of phenotypic and genotypic data were performed using the R software version 4.1.2 (R-Core-Team, 2020)

At each date of observation, phenotypic data were adjusted within each environment to correct for spatial environmental effects using information on row and column (Supplementary Method 2). After removing environments with insufficient phenotypic expression, we extracted Max and AUDPC (Supplementary Method 2). These variables were transformed using Box-Cox to correct for heteroscedasticity and non-normality of error terms (Supplementary Method 2). Variance analysis was performed using mixed linear models to quantify the variance associated to genetic, G×E, and non-additive effects (Supplementary Method 3 and Supplementary Method 4).

### Linkage disequilibrium

Intrachromosomic linkage disequilibrium (LD) was measured with the r^2^ value estimated by using VCFTools v0.1.16 [98]. The r^2^ was calculated for every combination of SNPs within a sliding window of 10 Mbp for peach and of 30 Kbp for apricot for every chromosome individually. We used a threshold of 0.2 to set the LD decay which was then represented graphically using Hill and Weir equation [99].

### Genome–wide association studies (GWAS)

Across-environment GWAS based on main additive and non-additive genetic as well as single-environment GWAS (Table 3) were performed using the single locus linear mixed model of [100] implemented in the R package MM4LMM [101]:

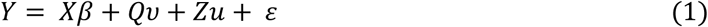

**Table 3.** Summary of the different GWAS model used in this study

| GWAS name | R Package | Rationale |
| --- | --- | --- |
| Main genetic effect additive GWAS | MM4LMM | Decipher the main additive genetic effects across environments |
| Main genetic effect non-additive GWAS | MM4LMM | Decipher the main non-additive genetic effect across environments |
| Single environment GWAS | MM4LMM | Study genetic effects environment by environment |
| Multi-environment GWAS | MTMM | Assess the effect of G × E interactions on QTL detection and identify genetic factors with common effects, *i.e.* stable marker effects across environments, or interaction effects, *i.e.* differential marker effects across environments |
| Meta-analysis GWAS | MetaGE | Assess the effect of G × E interactions on QTL detection by a joint analysis of single-environment GWAS results |

where *Y* is respectively the vector of G-BLUPs for across-environment GWAS or of environment-specific BLUPs for single-environment GWAS, *X* is the molecular marker score matrix, β is the vector of marker effects, *Q* is the structure matrix estimated by the SNMF R package [102] (Supplementary Method 5 and Supplementary Result 1), *v* is the fixed effect of the structure, *Z* is an incidence matrix, *u* is the vector of random background polygenic effects with variance σ^2^_u_ = K σ^2^_G_ (where K is the variance-covariance matrix of G, and σ^2^_G_ is the genetic variance), and ε is the vector of residuals.

The estimation of the variance-covariance matrix of G was determined by a genetic relatedness (or kinship) matrix, derived from all SNPs except those on the chromosome containing the SNP being tested as in [103]. According to the model, additive or non-additive kinships were used. They were calculated using the method developed by [104] and the source code published by [105]. For every pest or disease, we tested two different corrections including the kinship (model ‘K’) or both kinship and population structure (model ‘Q + K’). The best model was chosen on a trait-by-trait basis by comparing the likelihood of each null model (*i.e.* with only kinship of kinship + structure effects) using the AIC. We used a ‘modified’ 5% Bonferroni correction described by [106] to define significant associations between phenotypic data and genotypic markers. This threshold is based on the effective number of independent SNPs calculated thanks to the *Meff* equation implemented in the R package poolR [107]. The number of effective markers was 1326 for peach and 1778 for apricot.

To explore non-additive genotype–phenotype associations, we transformed the genotypic datasets following [108] and [54], as illustrated in Supplementary Table 4.

Subsequently, we performed single locus multi-environment GWAS by using the multi-trait mixed model (MTMM) as described by [29] in order to distinguish common and environment-specific marker–phenotype association (Table 3 and Supplementary Method 6). We finally performed a meta-analysis relying on the results of the single-environment GWAS model by using MetaGE R package [34] (Table 3 and Supplementary Method 7).

### Identification of high confidence QTLs and variance analysis of G×E and QTL×E

For all GWAS analyses, every QTL region was defined using the position of the top SNP (*i.e.* with the lowest *P*-value) and the estimated LD decay for every chromosome. In other words, QTLs were centred on the top SNP position with an extent of 2 time-specific chromosomal LD decay. Candidate SNPs distant less than the LD decays size were considered as belonging to this QTL. Each QTL name was defined as the associated trait name, followed by the chromosome number and the QTL’s order among all QTLs for this trait along the chromosome.

For every trait, we selected ‘high confidence’ QTLs which correspond to QTLs significant in at least 50% of the different additive GWAS model tested for the maximum of damage score or for AUDPC. We considered that each environment of the single-environment GWAS as separate models as well as the model ‘full’, ‘common’ and ‘gei’ of MTMM.

To assess the biological relevance of the high confident QTLs, we followed the approach previously described by [27,109] which aim at comparing the variance of a multi-locus multi-environment mixed model with and without the fixed effects of the QTLs and of the QTLs by environment interaction (Supplementary Method 8). Last, we calculated the indicator *γ_q,r_* to assess the proportion of genetic, G×L interaction and G×Y interaction variance explained jointly by all detected QTLs for each trait (Supplementary Method 8).

### Candidate gene analysis

Each high confidence QTLs region was queried in the *Prunus persica* Whole Genome v2.0 Assembly and Annotation v2.1 [39] or in the *Prunus armeniaca* Marouch whole genome v1.0 [41] from the Genome Database for Rosaceae (www.rosaceae.org). SNP gene localization and function were determined using the JBrowse tool (www.rosaceae.org/tools/jbrowse). Their function was then investigated in the literature, and genes involved in traits linked to host defence mechanisms were therefore proposed as putative candidate genes.

For peach, the local LD was calculated between the top SNP of the different ‘high confidence’ QTLs and all the SNPs belonging in this QTLs by using LDcorSV R package which takes into account population stratification and the relatedness between genotypes [110]. We used r^2^ > 0.2 to filter the putative candidate genes.

For genes which overlapped among several confidence QTLs for apricot, pairwise LD from all physically close SNPs in the region was computed using r^2^ calculated with LDcorSV R package [110]. The haplotypes were constructed by considering all top SNPs of the confidence QTLs involved for each G-BLUPs phenotype. Mean phenotypic distributions of the different haplotypes were compared using paired samples Wilcoxon tests for the associated G-BLUPs phenotypes.

## Acknowledgements and Funding

This work was supported by funding through LabEx AGRO 2011-LABX-002 (under I-Site Muse framework) coordinated by Agropolis Foundation (project ID: 2002-030), the INRAE Department for Plant Genetics and Breeding, the France AgriMer CASDAR Project ‘RésiDiv’ (project ID: 6846752) and the EU Horizon Innovation Actions InnOBreed n°101061028.

We are thankful to the two experimental research stations SEFRA and SICA-Centrex, to the INRAE experimental unit A2M and to the INRAE research experimental unit UERI of Gotheron for the maintenance of the two core collections. We thank Thierry Pascal for initiating the peach collection. We also thank Freddy Combe (INRAE, UERI Gotheron), Amandine Fleury (INRAE, UERI Gotheron), Luana Gillet (INRAE, UR GAFL), Béatrice Monnet (INRAE, UE A2M) as well as several students for their valuable contribution in field experiment and plant phenotyping. We are also grateful to Guillaume Roch (CEP Innovation) and Guy Clauzel (CEP Innovation) for grafting, pruning and DNA sampling the apricot core collection, to Jacques Lagnel (INRAE, UR GAFL) and Frédérique Bitton (INRAE, UR GAFL) for coordinating the bioinformatic processing of the raw data from the re-sequencing of the apricot core collection and to Océane Perez (INRAE, UR GAFL) for the preliminary analysis on apricot. We acknowledge Emilie Millet (INRAE, UR GAFL) for sharing her expertise in statistical analysis and providing valuable scientific advice.

## Author contributions

MR, BQ and JMA conceived the project and designed the experiment. JMA, BQ, AB and LB created the core collections. ND collected the DNA samples and coordinated the genotyping of the peach core collection. JMA collected the DNA samples and coordinated the resequencing of the apricot core collection. LH contributed to the alignment and filtration of the raw data from re-sequencing. MS, AB, LB, FG, MLP, MR, VSi and SV realised the phenotypic observations. MS analysed the data under the supervision of MR. VSe supervised GWAS analyses. LH contributed to the identification of candidate genes. MS wrote the manuscript. MR, BQ, JMA and VSe reviewed and edited the manuscript. All authors approved the manuscript.

## Data availability statement

The raw phenotypic and genotypic datasets and the R scripts generated for this study are accessible on the data.gouv public repository at: https://doi.org/10.57745/2HNRN0

